# Input-specific gating of NMDA amplification via HCN channels in mouse L2/3 pyramidal neurons

**DOI:** 10.1101/2023.09.25.559198

**Authors:** Viktor János Oláh, Jing Wu, Leonard K. Kaczmarek, Matthew JM Rowan

**Author notes:** Address for Correspondence: Matt Rowan, PhD, Emory University School of Medicine Department of Cell Biology, Atlanta, GA, 30322.

## Abstract

Layer 2/3 pyramidal cells (L2/3 PCs) play a crucial role in cortical information transfer. Although the dendritic arbors of L2/3 PCs are impressive, they often lack the distinct anatomical compartments characteristic of deeper L5 PCs. For example, many L2/3 PCs do not display an apparent distal tuft region. However, L2/3 PCs receive inputs from both thalamic (bottom-up) and cortical (top-down) inputs, which preferentially synapse onto their proximal and distal dendrites, respectively. Nonuniform organization of channels and NMDA receptors in L2/3 dendrites could serve to independently modulate these information streams to affect learning and behavior, yet whether L2/3 PC dendrites possess this capability has not been established. Here we found a previously unappreciated, non-uniform HCN channel distribution in L2/3 PCs, allowing for pathway-specific gating of NMDA receptor recruitment at bottom-up (proximal) but not top-down (distal) synapses. HCN availability shifted depending on developmental stage and neuromodulation, suggesting that the gain of thalamic and cortical-cortical signals in L2/3 may be independently modified *in vivo* across different timescales.

## INTRODUCTION

Layer 2/3 pyramidal cells (L2/3 PCs) are the most numerous excitatory neurons in the neocortex. As a major component of the canonical circuit, L2/3 PCs are thought to play crucial roles in information transfer and gain control of cortical output(1). Thus, disambiguating the afferent circuit connections of L2/3 PCs has been a significant focus of neuroscience research in the past decade(2–5). However, understanding cellular mechanisms by which L2/3 PCs transform information is equally important (6–9). Like their more extensively studied counterparts (e.g., L5 PT-type PCs, hippocampal CA1 PCs), L2/3 PCs are equipped with an array of dendritic nonlinearities(10, 11), enabling complex subcellular computations. Nonetheless, L2/3 PCs exhibit marked disparities from other PC types as well. Their morphology is less segmented due to the lack of a distinctive apical tuft (except for the deepest layer 3 cells (12)). Their dendritic protrusions are also shorter and consequently possess less surface area over which specialized nonlinear operations may be actualized. Therefore, the applicability of currently established neuronal dynamics to L2/3 PCs is questionable.

Dendritic voltage-gated channels and receptors are crucial for diversifying the computational power of neuronal cells, by allowing for nonlinear processing of arriving synaptic inputs(13). Nonlinear processing is essential for proper brain function and has been implicated in critical tasks such as conscious perception(14–16) and processing of sensory and motor input(17). Therefore, a more comprehensive understanding of subcellular signal processing dynamics in L2/3 PCs is warranted. Dendritic hyperpolarization-activated nonselective cation (HCN) channels are a staple of several PC classes(18, 19), playing essential roles in regulating resting membrane potential, the normalization of synaptic events arriving at spatially mismatched locations, and establishing oscillation frequency-selectivity(16). Beyond exerting control over the general excitability of neurons, HCN channels contribute to synaptic plasticity, and can regulate working memory(16). HCN dysfunction can also contribute to epilepsy and pain(20). Electrophysiological detection of this current often relies on identification of a characteristic ‘sag’ potential, elicited by hyperpolarizing current injections. In L2/3 PCs, this sag potential is negligible in most cortical areas (but see (21)) thus HCN channels are often presumed to be missing from mouse L2/3 PCs (22). However, it has recently been proposed that HCN is a key determinant of L2/3 PC passive properties at membrane potentials close to rest, despite the lack of traditional HCN channel physiological markers (23). In contrast to mice, the presence of HCN channels in L2/3 PCs of primates and humans is unambiguously supported by functional and anatomical evidence as well (24, 25) (26).

Here we report that L2/3 PCs throughout the cortex express functionally relevant HCN channels with unexpected functions. We found that HCN channels robustly influence the overall excitability of L2/3 PCs. Interestingly, we found that HCN channel availability in L2/3 PCs is biased in a unique proximal somato-dendritic manner, which is inverted with respect to L5 PT-type and hippocampal CA1 PCs (but see (27)), where HCN is enriched within their distal apical tufts(19, 28). In cooperation with local NMDA receptors, we found this L2/3 HCN organization could gate the amplification of proximal ‘bottom-up’ dendritic inputs, without affecting distal ‘top-down’ dendritic inputs. Furthermore, we found that HCN expression is robustly upregulated during early postnatal development, and that neuromodulation in adulthood can rapidly adjust HCN channel availability. Our results demonstrate that L2/3 PCs not only express dendritic HCN channels, but also employ these conductances in a previously unappreciated synaptic input-selective manner.

## RESULTS

### HCN channel expression in L2/3 PCs

HCN channels have been extensively documented and rigorously characterized in several cortical PC types(18), however, whether PCs residing in layer 2/3 of the mouse cortex express functionally relevant HCN channels remains ambiguous. Nonetheless, open-source snRNAseq (29) shows comparable HCN subunit mRNA levels in all cortical PC types of the visual cortex (Fig. 1a). Therefore, we investigated whether this mRNA signature translated to HCN surface expression. Immunohistochemical staining against HCN1 showed discernible somatic fluorescent signal in many cells residing in superficial layers of the cortex (Fig. 1b) along with staining of putative dendrites. To examine the potential functional role for HCN channels in L2/3 PCs, we performed *ex vivo* cell-attached recordings from visually identified L2/3 PCs in primary visual cortex slices from mature mice. Spike output upon extracellular stimulation (50 Hz) in layer 4 was then measured in control conditions and following pharmacological HCN channel block (Fig. 1c). We found that bath application of ZD-7288 (50 µM), a conventional blocker of HCN channels, significantly increased their firing (Fig. 1d,e). Similar results were obtained using another well-accepted HCN channel blocker; external Cs^+^ (2 mM, Fig. 1f,g). This effect was unlikely due to presynaptic modulation of these synapses, as the paired-pulse ratio was unchanged in a set of separate whole-cell recordings coupled with stimulation in layer 4 (Fig. 1h). Together, these results indicate that L2/3 PCs express a functionally relevant set of HCN channels which can modulate AP firing following synaptic stimulation.

**Fig 1.**
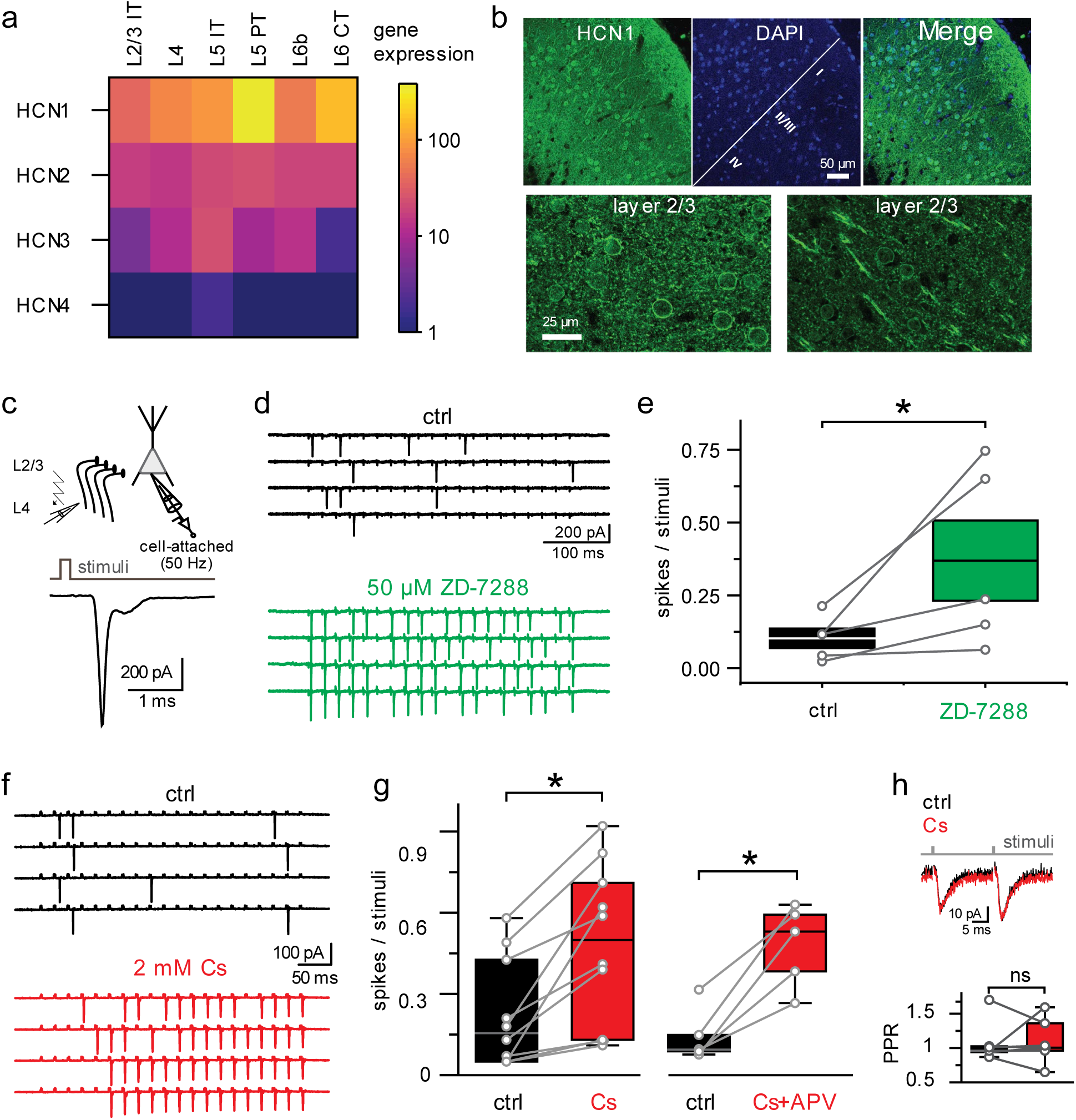
HCN expression and effect of Cs^+^ on AP firing in cortical L2/3 PCs. **a.** HCN gene subunits show uniform expression pattern among cortical PC types. Data is collected from the online available dataset(29). **b.** HCN1 immunohistochemistry reveals dense somatic expression in layer 2/3 of the visual cortex **c.** HCN channels control the excitability of L2/3 PCs. Schematic illustration of the experimental arrangement of cell attached recordings during extracellular stimulation of L4, and the shape of the elicited action potential. **d.** Representative traces showing increased firing upon ZD-7288 application. **e.** ZD-7288 increases the firing of L2/3 PCs upon electrical stimulation (0.1 ± 0.03 spikes/stimuli vs 0.4 ± 0.12 spikes/stimuli in control vs ZD-7288 application, respectively, p=0.04, t(4)=-3.0, Student’s paired-sample t-test, n=5). **f.** Increased spiking of L2/3 PC upon blockade of HCN channels with 2 mM CsMeSO4 (right). **g.** Pharmacological block of I_h_ significantly increases the firing response of L2/3 PCs (0.22 ± 0.06 spikes/stimuli vs. 0.48 ± 0.09 spikes/stimuli for control vs CsMeSO4 bath application, p=0.002, t(9)=-4.42, Student’s paired t-test, n=10. **h.** Presynaptic contributions were examined using the paired-pulse ratio (PPR) in whole cell current clamp recordings (1.09 ± 0.12 spikes/stimuli vs. 1.17 ± 0.15 spikes/stimuli for control vs CsMeSO4 bath application, p=0.422, t(6)=-0.8612, Student’s paired t-test, n=7).

### HCN channels constrain the overall excitability of L2/3 PCs

To investigate the role of HCN channels in regulating the excitability of L2/3 PCs, we performed further L2/3 PC whole-cell recordings. A conventionally used assessment of HCN channel function is the characterization of the sag potential(18). This hallmark of I_h_ activation is measured as the amount of rectification detected in response to hyperpolarizing current injections. In line with previous reports(30), we found that L2/3 PCs exhibited an unremarkable amount of sag potential at ‘typical’ current commands. However, upon larger negative current injections, a discernable rectification was revealed (Fig. 2a). In addition, the rectification of the steady-state responses was abolished in the presence of external Cs^+^, which could be restored by artificial I_h_ supplementation by dynamic clamp (Fig. 2b). Similarly, sag became non-detectable when I_h_ was blocked, which could then be online restored during recordings by dynamic clamp (Fig. 2c,d). Furthermore, we found that passive electrical properties were also significantly modulated by HCN blockade (Fig. 2e,f, Supplementary Fig. 1), however, action potential halfwidth was unaffected (1.04 ± 0.09 ms vs. 1.12 ± 0.17 ms for control vs. CsMeSO4 bath application; p=0.324, t(6)=-1.073, Student’s paired t-test; n=7. Together, these results explain the dramatic effect of HCN blockade on L2/3 excitability (Fig. 1). The non-apparent sag potential at conventional hyperpolarizing current steps is thus surprising, suggesting a markedly faster occurrence of the I_h_ effect exists in L2/3 PCs which occludes the biphasic nature of the sag response. To investigate this possibility, we measured the time constant of the immediate voltage responses. We found a strong positive correlation between the amplitude of a passive current command and the voltage response time constant, which was abolished by Cs^+^ bath application (Fig. 2g). This confirms that I_h_ has a strong effect during the initial phase of hyperpolarizing voltage responses, which may contribute to the lack of detectable sag potential at modest current injections. However, other biophysical mechanisms may also contribute to the muted sag effect in L2/3 PCs.

**Fig 2.**
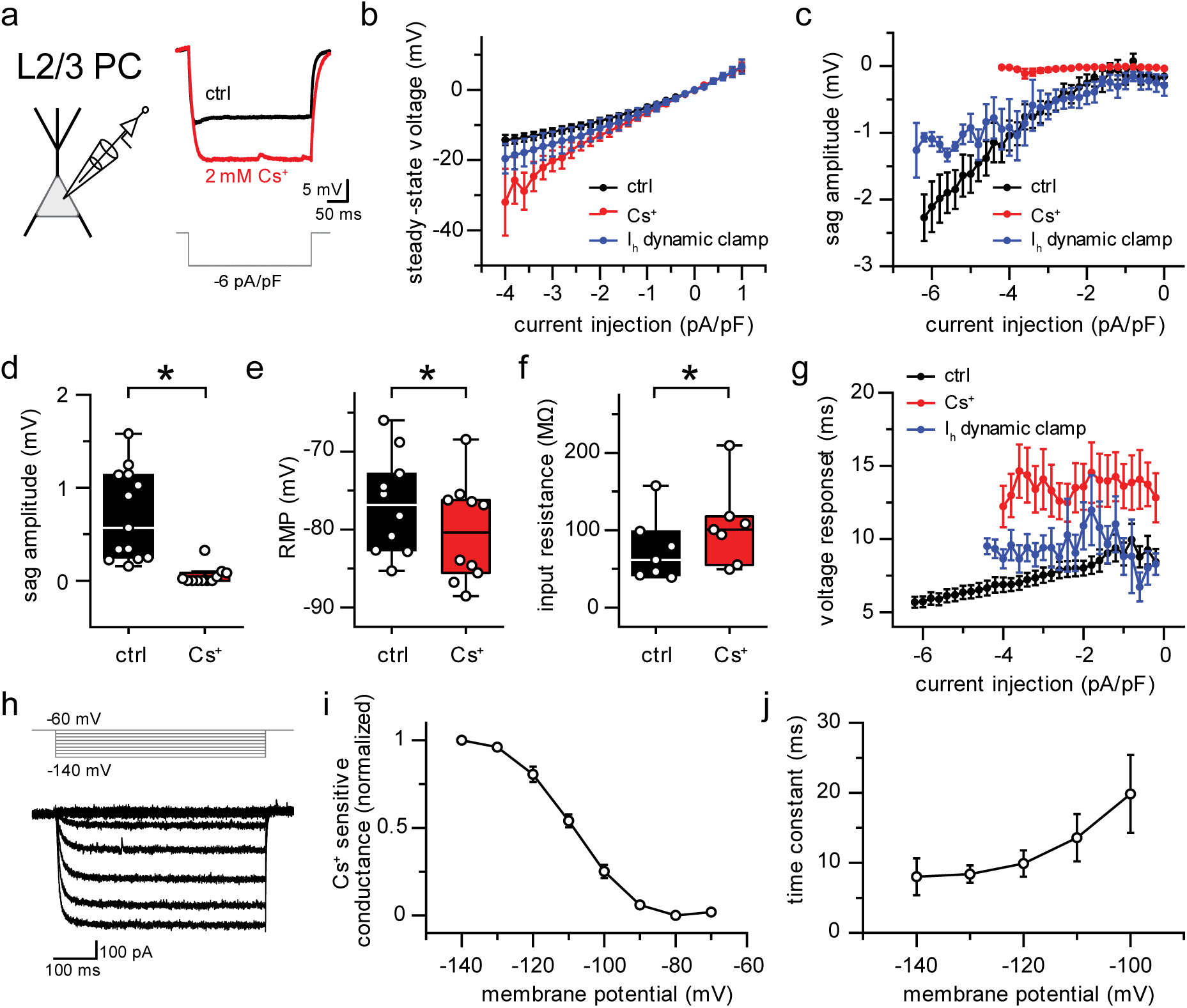
Functionally relevant I_h_ in L2/3 PCs. **a.** Whole cell current clamp recordings showing the presence and absence of sag response to very negative current commands in control conditions (black) and upon Cs^+^ application (red). **b.** Steady-state voltage response rectification upon hyperpolarizing current injections is mediated by I_h_. **c.** Sag voltage is abolished by blocking I_h_. **d.** Sag amplitude is Cs^+^ sensitive (0.7 ± 0.49 mV vs. 0.05 ± 0.1 mV for control vs CsMeSO4 application, p=2.46^-4^, t(22)=4.37, Student’s two-sample t-test, n=13 and 11). **e.** Resting membrane potential is modulated by I_h_ (-76.75 ± 6.37 mV vs. -80.29 ± 6.48 mV for control vs CsMeSO4 bath application, p=1.98^-4^, t(9)=6.02, Student’s paired t-test, n=10). **f.** Input resistance is altered by I_h_ (74.78 ± 42.34 mV vs. 104.99 ± 52.91 mV for control vs CsMeSO4 bath application, p=0.048, t(6)=-2.48, Student’s paired t-test, n=7). **g.** The time course of voltage responses to hyperpolarizing current on the magnitude of the current injection is abolished by Cs^+^. **h.** Example voltage clamp recording of cesium sensitive currents in a L2/3 PC. **i.** Voltage dependence of cesium sensitive current activation, **j.** Voltage dependent activation kinetics of cesium sensitive currents.

Next, we examined the voltage dependence and kinetic properties of I_h_ in L2/3 PCs in voltage clamp (Fig. 2h). Significant Cs^+^-sensitive (1mM) currents were activated at subthreshold membrane potentials, with half-activation voltages more negative than the resting membrane potential, and with relatively fast activation kinetics (Fig. 2i,j). Overall the pharmacological profile, effects on passive electrical properties, and voltage dependence all point to previously published data describing HCN channels (31). Together these results indicate that L2/3 PCs express a functionally relevant population of HCN channels, which constrain the excitability of these cells. Similar responses were observed in voltage clamp recordings across several cortical regions (e.g., in S1, M1, and A1; Supplementary Fig. 2), suggesting that HCN channels are a consistent feature in L2/3 PCs.

### Noncanonical subthreshold signal processing in L2/3 PCs by I_h_

Electrical signals at different dendritic compartments undergo distance-dependent distortion (28, 32), which has a major effect on neuronal function in cells with extensive dendritic branching (33). To counteract these effects, larger L5 PT-type PCs and CA1 PCs express HCN channels with nonuniform distributions (19, 31). As L2/3 PCs have a much more compact morphology, synaptic input distortions might be less severe, raising the possibility of an alternative function of HCN channels in these cells. Thus, to investigate the distance-dependence of HCN on L2/3 synaptic inputs, we made whole-cell patch recordings from Alexa-594 filled L2/3 PCs and performed extracellular minimal stimulation at different dendritic locations guided with 2P microscopy (Fig. 3a). Elicited EPSPs likely reflect the activation of a single axon, as stimulus intensity was set to achieve an ∼50% failure rate(34). As HCN deactivation has been shown to strongly impact the duration of synaptic events(28, 32), we quantified changes in the half-width of locally evoked EPSPs, here using the HCN-selective blocker ZD-7288 (50 μM). Surprisingly, our results were in contrast to previous reports in L5 PT-type and CA1 PCs, in that I_h_ block had a far smaller effect on the time-course of distally-evoked (>100 μm distance from the soma) synaptic responses. In contrast, proximal inputs (<100 μm) were markedly more broadened after I_h_ blockade (Fig. 3b,c). These results suggest that L2/3 PCs are equipped with a distinctive subcellular HCN distribution, potentially contributing to noncanonical distance-dependent input amplification along the dendritic axis of L2/3 cells.

**Fig 3.**
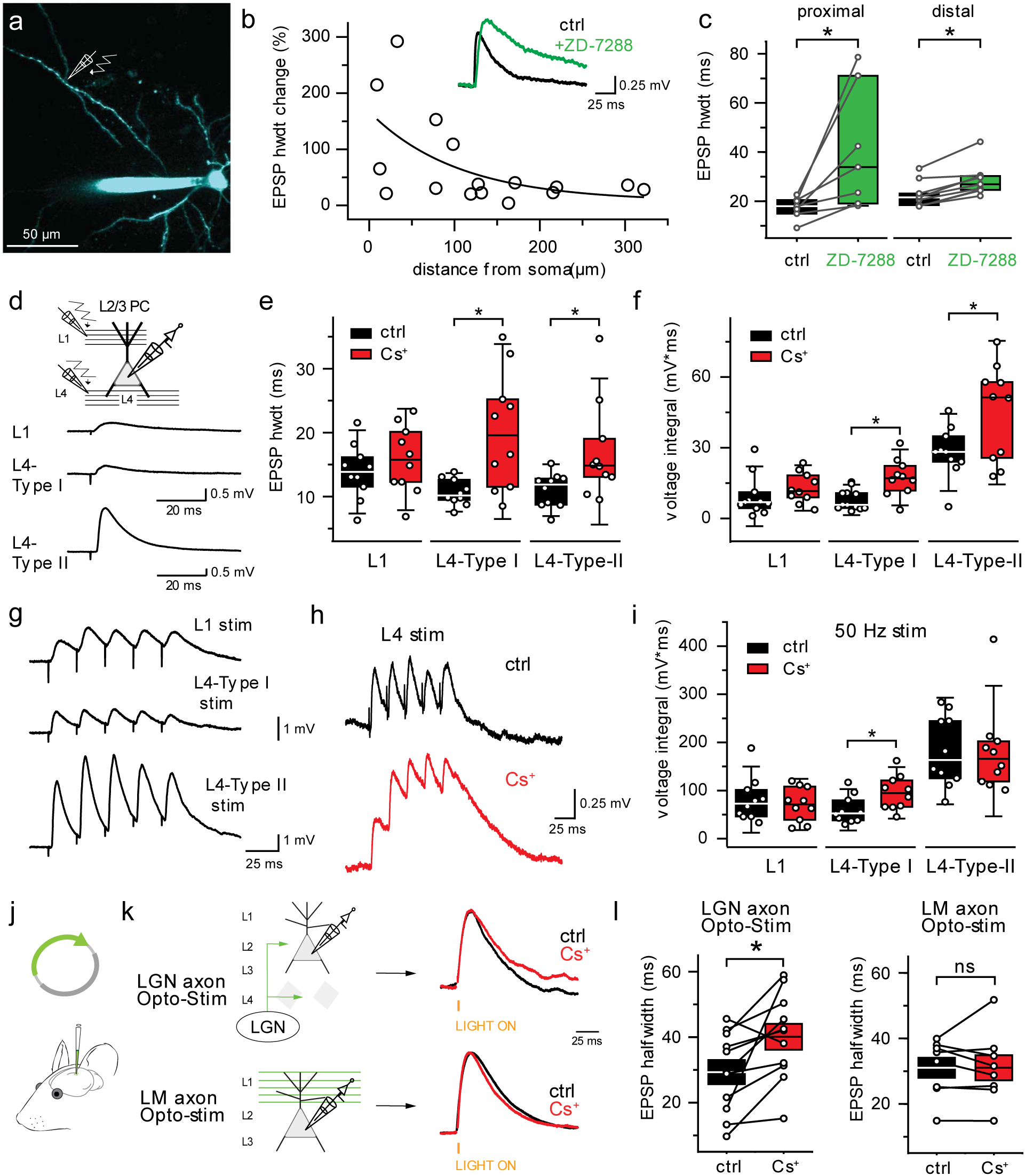
Noncanonical effect of I_h_ block on EPSPs in L2/3 PCs. **a.** Experimental arrangement of extracellular targeted stimulation of a L2/3 PC dendrite. **b.** Application of ZD-7288 revealed a proximal bias for EPSP modulation by I_h_ (Exponential fit τ=104.7 µm, R^2^=0.33, n=18). Inset shows a stimulating location 28 µm away from the soma in control conditions (black trace) and upon Z-7288 application (green trace). **c.** EPSP halfwidth is significantly modulated along the dendritic axis by I_h_, (proximal dendritic locations: 17.42 ± 1.68 ms vs. 40.95 ± 24.78 ms for control vs 50µm ZD-7288 bath application, p=0.03, t(6)=2.84, Student’s paired t-test, n=7, distal dendritic locations: 22.68 ± 1.78 ms vs. 28.65 ± 2.17 ms for control vs 50µm ZD-7288 bath application, p=2.1^-4^, t(8)=-6.39, Student’s paired t-test, n=9)**. d.** Schematic illustration of the experimental arrangement. L2/3 PCs were recorded in whole-cell current clamp mode, and a stimulating electrode was placed either in L1 or L4. **e.** I_h_ modulates EPSP halfwidth for L4 stimulation, but not for L1 (13.87 ± 1.37 ms vs. 15.8 ± 1.67 ms for control vs CsMeSO4 bath application in L1, p=0.38, t(18)=-0.89, Student’s two- sample t-test, n=10 each, 10.65 ± 0.64 ms vs. 20.17 ± 2.88 ms for control vs CsMeSO4 bath application for L4-Type I inputs, p=0.005, t(18)=-3.22, Student’s two-sample t-test, n=10 each, 11.01 ± 0.85 ms vs. 17.05 ± 2.41 ms for control vs CsMeSO4 bath application for L4-Type II, p=0.03, t(18)=-2.36, Student’s two-sample t-test, n=10 each). **f.** Pathway-specific I_h_ effect on unitary EPSP voltage integral (9.35 ± 2.54 mV*ms vs. 12.9 ± 2.04 mV*ms for control vs CsMeSO4 bath application in L1, p=0.3, t(19)=-1.07, Student’s two-sample t-test, n=11 and n=10 respectively, 7.85 ± 1.24 mV*ms vs. 17.39 ± 2.5 mV*ms for control vs CsMeSO4 bath application for L4-Type I inputs, p=0.002, t(20)=-3.6, Student’s two-sample t-test, n=12 and n=10 respectively, 28.01 ± 3.45 mV*ms vs. 44.89 ± 6.42 mV*ms for control vs CsMeSO4 bath application for L4-Type II inputs, p=0.03, t(18)=-2.31, Student’s two-sample t-test, n=10 each). **g.** Representative recordings of 50 Hz repeated extracellular stimuli (L1 – grey, L4-Type I – blue, putative L4-Type II -green). **h.** Temporal summation of L4-Type I input stimulation in control conditions (black) and during CsMeSO4 application (red). **i.** Pathway-specific I_h_ modulation of synaptic summation (81.05 ± 14.5 mV*ms vs. 71.07 ± 11.19 mV*ms for control vs CsMeSO4 bath application in L1, p=0.6, t(18)=0.54, Student’s two-sample t-test, n=10 each, 60.06 ± 9.03 mV*ms vs. 95.15 ± 11.29 mV*ms for control vs CsMeSO4 bath application for L4-Type I, p=0.03, t(18)=-2.42, Student’s two-sample t-test, n=10 each, 182.08 ± 23.36 mV*ms vs. 182.05 ± 28.59 mV*ms for control vs CsMeSO4 bath application for L4-Type II, p=0.99, t(18)=8.74*10^-4^, Student’s two-sample t-test, n=10 each). **j.** Schematic illustration of the experimental setup. Mice were injected with AAV.CAMKII.C1V1/eYFP either in the latermodial visual area (LM, to label axons arriving to L1) or in the LGN (to label proximal targeting axons). **k.** Example trace showing optogenetic activation of LM (top) and LGN (bottom) axons in control conditions (black) and upon Cs^+^ application (red). Traces are normalized. Blue line marks the timing of the optogenetic stimulus. **l.** Cs^+^ effect on optogenetically elicited EPSPs from cortico-cortical (LM, left; 35.06 ± 3.06 vs. 31.04 ± 3.82 for control vs Cs^+^ respectively, p=0.99, t(7)=0.001, Student’s paired-sample t-test, n=8) and thalamic (LGN, right; 29.31 ± 3.68 vs. 40.11 ± 3.95 for control vs Cs^+^ respectively, p=0.01, t(10)=- 2.95, Student’s paired-sample t-test, n=11) sources.

### Pathway-specific modulation of synaptic information

L2/3 PCs receive information from distinct brain regions, such as “top-down” (cortical-cortical) axons, generally arriving to layer 1(35) and “bottom-up” (thalamocortical/layer 4) axons, terminating on proximal dendrites located in layer 4 and 2/3 (36). Due to the stronger proximal effect of HCN on L2/3 PC EPSPs, we explored whether this modulation yielded input-pathway-specific differences. Whole-cell recordings were again obtained from L2/3 PCs, and EPSPs were elicited by minimal stimulation with electrodes situated to preferentially stimulate either layer 4 or layer 1 axons (electrode placement ∼200μm laterally from recorded PC in either layer). Inhibition was left intact in these experiments. While most evoked EPSPs were small (< 0.5 mV), a subset evoked with minimal stimulation was outsized (we deemed these here as L4-Type I and L4-Type II inputs, respectively; Fig. 3d, Supplementary Figure 3). The L4-Type II events were substantially larger than previous reports(3) but see (37).

I_h_ blockade significantly broadened elicited EPSPs when the stimulation electrode was placed in layer 4, but not in layer 1 (Fig. 3e). Consequently, the effect of Cs^+^ on the EPSP integral was significant for L4 inputs, but not L1 inputs (Fig. 3f). The effect of Cs^+^ on EPSP half-width and voltage integral was similar for both L4-Type I and L4-Type II inputs, suggesting intermingled termination zones(36) in proximal dendrites. HCN channels are notable for their ability to restrain temporal integration of synaptic inputs(28). Therefore, we repeated these experiments using more high-frequency stimulation (5 stimuli at 50 Hz; Fig. 3g; inhibition was blocked due to strong inhibitory recruitment at 50Hz). In accordance with the effect on single EPSPs, summation of L1 (i.e., distal) inputs was unaltered with HCN block. In contrast, HCN blockade led to increased temporal summation of L4-Type I inputs (Fig. 3h,i). Summation of L4-Type II inputs was not affected by HCN channel block, suggesting further diversification between L4-Type I and L4-Type II inputs as well.

To more accurately evaluate whether synaptic signaling from *bone fide* bottom-up and top-down axons are differentially regulated by HCN, we next injected a CaMKII-driven, C1V1-expressing AAV into either the LGN or the lateromedial visual (LM) areas in separate experiments (LM, Fig. 3j,k). LGN and LM-originating axons are thought to preferentially synapse onto proximal and distal L2/3 dendrites, respectively (38–41). In these experiments, short LED-directed amber light pulses could reliably elicit EPSPs in L2/3 PCs (Fig. 3k). We used a ‘minimal’ opto stim approach for these experiments, meaning we used as low light power as possible to reliability (1 ms pulses at 0.2 Hz) evoke EPSPs, while avoiding AP firing and thus further network recruitment. Inhibition was not blocked during optical stimulation. These optogenetic experiments revealed a negligible effect of Cs^+^ on top-down, LM-driven EPSPs. Conversely EPSPs were significantly broadened following Cs^2+^ of bottom-up, LGN-originating EPSPs (Fig. 3k,l). Together these results suggest that HCN channels preferentially regulate thalamic-originating, bottom-up nonlinear signaling in proximal dendritic regions of L2/3 PCs.

### I_h_ channels constrain NMDA receptor-dependent synaptic boosting in proximal dendrites

In a passive environment, dendritic filtering(42) creates attenuated (i.e., wider) EPSPs when originating further from the soma. Yet our results show an opposite phenomenon with HCN channels blocked. It has been described that reduced HCN availability can facilitate dendritic boosting mechanisms via activation of either voltage-gated(43) or NMDA(44) channels. To investigate possible contributions of dendritic voltage-dependent processes, we next performed NEURON computational simulations using a realistic L2/3 PC dendritic morphology(45). Inserting voltage-gated sodium or potassium channels with various distributions could not recapitulate our distance-dependent EPSP observations (Supplementary Figure 4.). Thus, we next investigated whether NMDA receptors might play a role in the observed phenomenon due to their described overwhelming contribution to *in vivo* activity in L2/3 PCs (8, 46).

We first tested whether NMDA receptors contribute to spontaneously occurring synaptic inputs (sEPSPs) in L2/3 PCs(47) in slice experiments. Due to potential alterations in spontaneous release and overall circuit activity, extracellularly applied NMDA blockers may introduce significant confounds for this line of investigation especially as spontaneous synaptic potentials show tremendous variability in frequency, amplitude, and kinetics. To overcome this conundrum, we designed a custom-built pipette attachment through which the intracellular solution could be stably exchanged during whole-cell recordings (48). We recorded sEPSPs in L2/3 PCs first using control intracellular solution, which was then exchanged to a 50 µM MK-801 (non-competitive NMDA receptor antagonist) containing intracellular solution (Fig. 4a). To also capitalize on the voltage-sensitivity of NMDA receptors, average peak amplitudes were recorded at two different membrane potentials for each intracellular condition; -80 mV and -55 mV (resting membrane potential, and a somatic membrane potential depolarized enough to partially unblock NMDA receptors (49)). Due to the small nature of L2/3 PC sEPSPs, quantification of other waveform parameters (i.e., EPSP width) was avoided. After quantifying the mean sEPSP peak at -80 mV, we calculated the expected (linear) sEPSP peak at -55 mV and compared this prediction to our observations of -55 mV. We found that sEPSPs were significantly larger than predicted after depolarization (Fig. 4b), alluding to the presence of a nonlinear subthreshold amplifying mechanism. We found that intracellular MK-801 solution exchange could severely dampen this synaptic amplification (Fig. 4c), indicating that NMDA receptors actively contribute to shaping naturally occurring sEPSPs in L2/3 PCs. The effect of pipette perfusion itself on our measurements was negligible (Supplementary Fig. 5)

**Fig 4.**
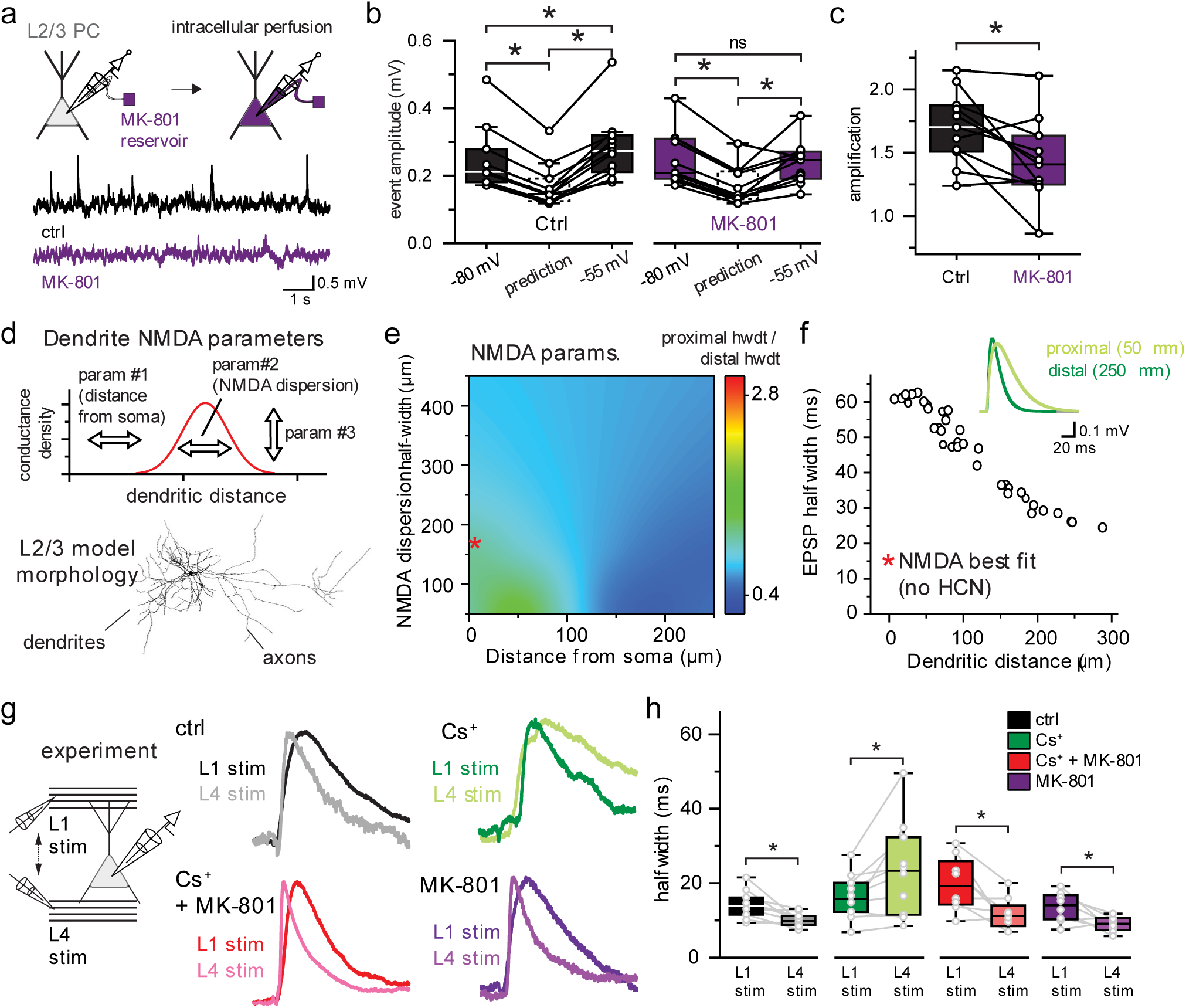
Pathway-specific NMDA receptor boosting of synaptic information. **a.** Schematic illustration of intracellular pipette solution perfusion during whole-cell patch clamp recordings. Briefly, the pipette was filled with control recording solution (black trace), which was exchanged to an intracellular solution containing 50 µM MK-801 (purple trace) during the recording. **b.** Mean peak amplitude of sEPSPs measured at two different membrane potentials and two different recording conditions (ctrl -80 mV vs. ctrl prediction: p=4.72*10^-6^, t(10) = 8.86, ctrl -80 mV vs ctrl -55 mV: p=0.02, t(10) = -2.81, ctrl prediction vs ctrl -55 mV: p=1.63*10^-5^, t(10) = -7.71, MK-801 -80 mV vs. MK-801 prediction: p=1.15*10^-6^, t(10) = 10.36, MK-801 -80 mV vs MK-801 -55 mV: p=0.58, t(10) = 0.57, MK-801 prediction vs MK-801 -55 mV: p=2.69*10^-3^, t(10) = -3.95, Student’s paired-sample t-test, n = 11 for each). **c.** Synaptic amplification estimated by dividing the predicted peak amplitude from the hyperpolarized recordings and the measured peak at depolarized membrane potentials (1.69 ± 0.09 and 1.43 ± 0.1 for ctrl and MK-801 amplification respectively, p=0.01, t(10) = 3.04, Student’s paired-sample t-test, n=11). **d.** Schematic illustration of the multiparameter NEURON fitting procedure for dendritic NMDA. Conductance localization, dispersions, and conductance densities of NMDA were varied along a somato-dendritic axis as set by a Gaussian curve, which was shifted in space, in peak, and in width (top panel). Simulations were carried out using fully reconstructed L2/3 PC from the Allen Institute open-source database (bottom panel). **e.** Inserting NMDA receptors into proximally located simulated synapses recreated the experimentally determined distance-halfwidth relationship. Notice the best fit of the control conditions (where proximal/distal EPSP halfwidth is similar), is characterized by a Gaussian curve with proximal peak center with a moderate width (dispersion). **f.** Best fit of the distance-dependent EPSP halfwidth with no HCN in the system. Red asterisk denotes the best fit parameter-space on panel. Proximally (gray) and distally (black) originating EPSPs from the model are shown in the inset. **e. g.** Amplitude-normalized EPSPs recorded from L2/3 PCs by minimal stimulation in either in L1 (dark color tones) or L4 (light color tones) in different recording conditions (ctrl: *black*, extracellular cesium: *green*, extracellular cesium + intracellular MK-801: *red*, intracellular MK-801: *purple*). Cells were recorded for each pharmacological condition independently with L4 and L1 stimulation occurring in each recording. **h.** As predicted (panel f) proximal inputs from L4-originating EPSPs were faster than those from distal L1 synapses in control conditions. The broadening effect of cesium on L4 (proximal) originating EPSPs is occluded by NMDA receptor block (14.25 ± 1.17 ms and 10.11 ± 0.56 ms for L1 vs L4 stimuli in control conditions, p=6.73*10^-3^, t(9)=3.5, Student’s paired-sample t-test, n=10 each, 16.22 ± 1.91 ms and 23.61 ± 4.04 ms for L1 vs L4 stimuli during Cs bath application, p=0.03, t(9)=-2.67, Student’s paired-sample t-test, n=10 each, 19.92 ± 2.65 ms and 11.8 ± 1.55 ms for L1 vs L4 stimuli during Cs bath application and MK-801 intracellular application, p=0.02, t(7)=3.01, Student’s paired-sample t-test, n=8 each, 13.62 ± 1.43 ms and 8.97 ± 0.72 ms for L1 vs L4 stimuli during MK-801 intracellular application, p=0.025, t(7)=2.84, Student’s paired-sample t-test, n=8 each).

Having established that NMDA receptors play an active role in modulating sEPSPs, we next utilized computational simulations to explore whether non-uniform NMDA recruitment could explain the subthreshold boosting in proximal L2/3 synapses, as described earlier following HCN channel blockade (Fig. 3b). Accordingly, this initial model did not include HCN channels. NMDA was inserted into a NEURON model with varying distributions and conductance densities (Fig 4d) to systematically evaluate a large parameter space. These simulations (Fig. 4e) revealed that a proximally-biased NMDA distribution (50) could indeed produce local EPSP boosting (Fig. 4e,f) similar to our experimental findings in an HCN-lacking condition.

Because HCN channels can reduce NMDA recruitment during synaptic signaling, we next explored whether proximal enrichment of HCN allows for bottom-up-specific gating of NMDA amplification (Fig. 4e,f) now using complimentary slice experiments. To accomplish this, we recorded L2/3 EPSPs originating from both proximal L4 and distal L1 inputs during each recording (i.e., two stimulation locations for each cell; Fig. 4g). For each recording, one pharmacological condition was evaluated to sequentially explore the interplay between HCN and NMDA (Fig. 4g). In control conditions, L4-evoked EPSPs were slightly narrower than L1-evoked EPSPs. This layer-specific relationship was inverted following extracellular Cs^+^ to block HCN (Fig. 4g,h). This result recapitulated our NEURON simulation best-fit prediction, where a proximally-biased NMDA distribution was necessary (Fig. 4f).

To confirm that this L4-specific EPSP broadening after HCN channel block was indeed due to NMDA-inducted boosting, as predicted in our simulations (Fig. 4f), we next examined L4 and L1- evoked EPSPs using Cs^+^ and intracellular MK-801. In the presence of internal MK-801, the broadening effect of Cs^+^ on L4-evoked EPSPs was abolished, thereby reverting the L4-L1 EPSP relationship to a control-like state. (Fig. 4g,h). The NMDA receptor contribution is unlikely related to different event amplitudes in proximal and distal sources, as the peak and halfwidth of the elicited EPSPs in our dataset showed no correlation (Supplementary Fig. 6). Together, our modeling and experimental results indicate that NMDA-directed amplification of L4 inputs is strongly constrained by a proximal HCN bias in L2/3 PCs dendrites, allowing for pathway-specific modulation of ‘bottom-up’ inputs without affecting those arriving from ‘top-down’ L1 axons.

### Noncanonical HCN distribution predicted in L2/3 PCs

We next carried out a more complete computational simulation which included both HCN and NMDA in a L2/3 PC. To test whether our results could be explained due to a non-uniform HCN distribution, HCN channels were inserted into the model with varying distributions and conductance densities (Fig 5a) to evaluate a large parameter space. Our simulations revealed that proximal bias in HCN channel distributions could best recapitulate our observations of proximally and distally originating EPSP waveforms in control conditions (Fig. 5b,c). This is in line with the proximal HCN1 channel staining (Fig. 1) revealed by immunohistochemistry. Together our results indicate that L2/3 PCs have a proximal HCN channel bias, whereas previous studies found a distal dendritic HCN bias in L5 PT-type PCs and CA1 PCs(19, 31), which could constrain NMDA receptor amplification in an input-specific manner.

**Fig 5.**
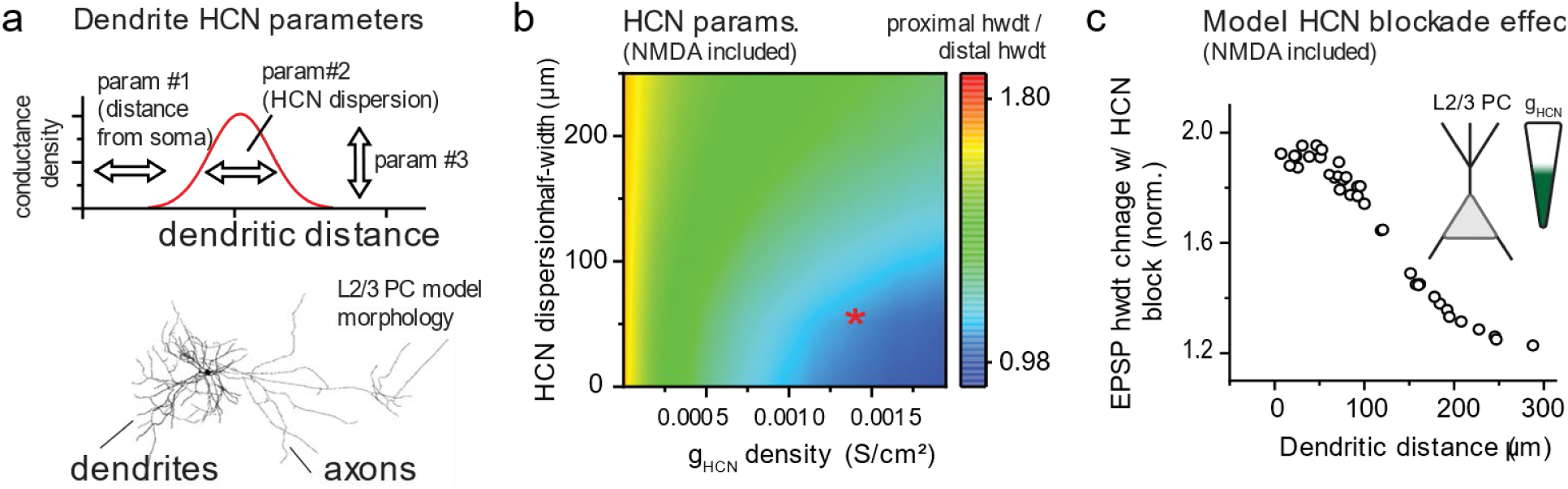
Proximal bias in HCN channel expression dampens proximal NMDA receptor-dependent synaptic boosting in a NEURON model simulation. **a.** Schematic illustration of the multiparameter NEURON fitting procedure for dendritic HCN. Simulations here included the NMDA dendritic parameters established in Fig. 4. HCN conductance localization, dispersions, and conductance densities were varied along the somato-dendritic axis as set by a Gaussian curve, which was shifted in space, in peak, and in width as well (top panel). Simulations were carried out using fully reconstructed L2/3 PC from the Allen Institute open-source database (bottom panel). **b**. A proximal abundance bias of HCN channels recapitulates experimental findings in control conditions. Best fit in the parameter space tested is denoted with red asterisk, where a high density of HCN channels were proximally localized. **c.** Model-derived data showing the effect of HCN blockade in the best-fit scheme when NMDA is also included as shown in Fig. 4. Note that the effect of HCN removal in this model closely recapitulates our experimental findings using ZD-7288 earlier.

### Tight developmental upregulation of Layer 2/3 HCN channels

Due to the ability of HCN channels to regulate NMDA boosting shown earlier, modifying HCN availability or expression may represent an important activity or plasticity gate in L2/3. Previously we found that HCN recruitment was upregulated in interneurons during early development(51). Indeed, several ion channels show postnatal maturation in their biophysical properties and surface expression(52), raising the possibility of a different proximo-distal HCN bias in young L2/3 PCs compared to adult animals. To examine this possibility, L2/3 PCs were recorded at three developmental time points (1^st^, 2^nd^ and 6^th^ postnatal week) in S1 cortex. Our assessment showed that a rapid developmental HCN upregulation occurs between 1 and 2 weeks old. We also found that 1 week old L2/3 PCs displayed markedly different passive properties, while 2-week-old cells were quite similar to mature cells (Fig. 6a). The non-quantifiable sag potential in 1 week old animals implies the lack of I_h_ current at this developmental time point, however, the disproportionately large input resistance and elevated resting membrane potential also suggests that other background leak channels may also be missing at this developmental window (Fig. 6b). Subsequent voltage clamp recordings confirmed that in 1 week old animals, L2/3 PCs do not possess a considerable amount of I_h_ in contrast to 2 and 6 week old animals (Fig. 6c,d). We found that both the resting membrane potential and peak I_h_ conductance changes throughout development were tightly (negatively) correlated (Fig. 6e). This provides additional evidence for parallel maturation of HCN with other leaky channels, as HCN channel expression alone would be predicted to shift the membrane to more depolarized potentials (Fig. 2e).

**Fig 6.**
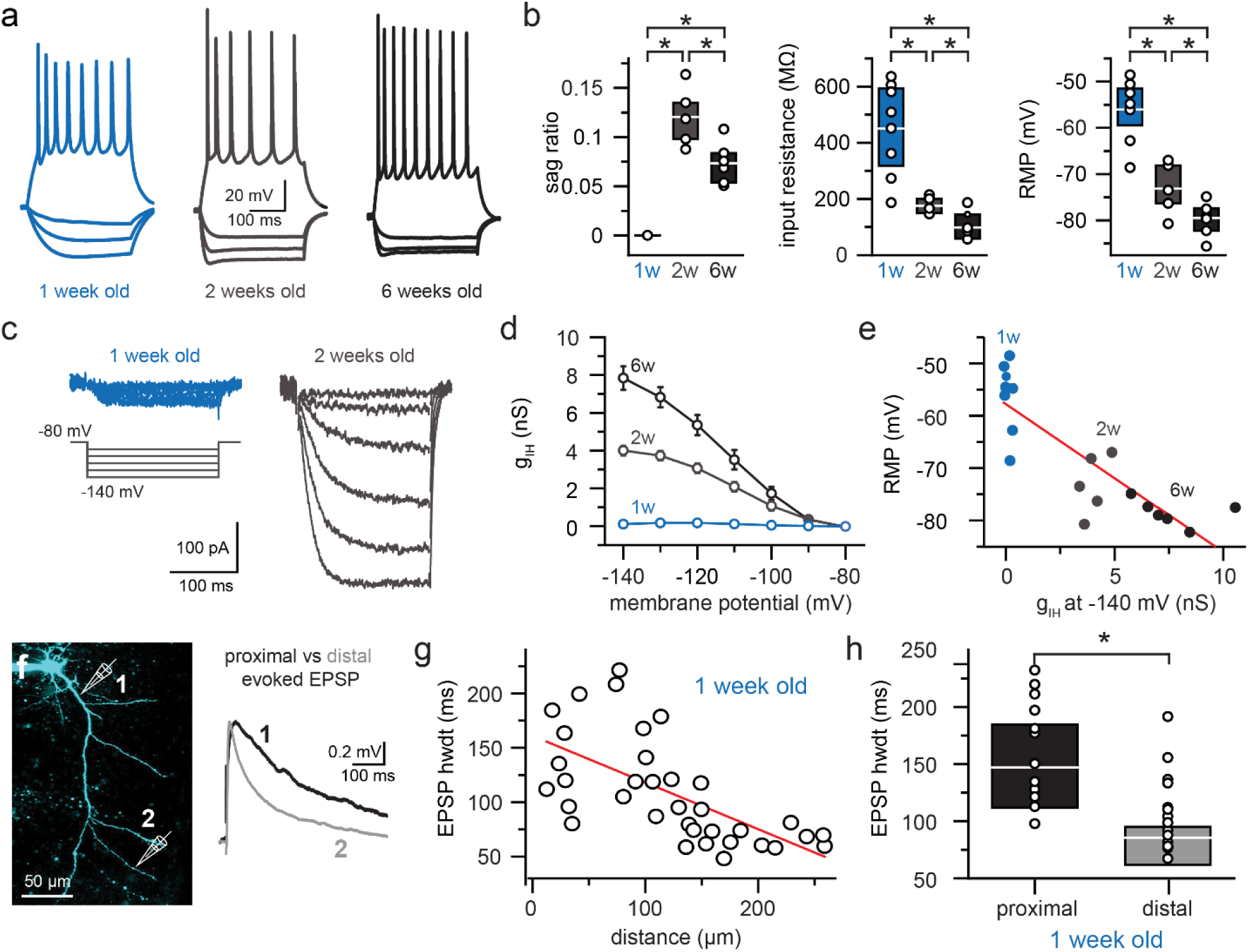
Developmental regulation of I_h_ expression. **a.** Representative firing patterns recorded from one week old (left, blue), two weeks old (middle, grey) and six weeks old L2/3 PCs. **b.** Developmental regulation of sag ratio (0.0 ± 0.0, 0.12 ± 0.01 and 0.07 ± 0.01 for one- (n=8), two- (n=5) and six week old L2/3 PCs (n=6), respectively, p=0 for one week old vs two weeks old cells, p=0.01, t(9)=3.04 for two vs six weeks old cells and p=0 for one vs six weeks old cells, Student’s two-sample t-test), input resistance (450.6 ± 57.91 MΩ, 175.66 ± 13.66 MΩ and 98.34 ± 18.86 MΩ for one- (n=8), two- (n=5) and six week old L2/3 PCs (n=7), respectively, p=0.004, t(11)=3.65 for one week old vs two weeks old cells, p=0.01, t(10)=3.06 for two vs six weeks old cells and p=1.11*10^-4^, t(13)=5.54 for one vs six weeks old cells, Student’s two-sample t-test) and resting membrane potential (-56.05 ± 2.33 mV, -73.13 ± 2.55 mV and -79.47 ± 1.33 MΩ for one- (n=8), two- (n=5) and six week old L2/3 PCs (n=7), respectively, p=5.9*10^-4^, t(11)=4.76 for one week old vs two weeks old cells, p=0.04, t(10)=2.39 for two vs six weeks old cells and p=1.36*10^-6^, t(13)=8.37 for one vs six weeks old cells, Student’s two-sample t-test). **c.** Representative voltage clamp recordings of hyperpolarization activated currents in one week old (left, blue), two weeks old (middle, gray) and six weeks old L2/3 PCs (right, black). **d.** Voltage dependence of hyperpolarization activated currents in one week old (blue), two weeks old (gray) and six weeks old L2/3 PCs (black). **e.** Negative correlation between measured hyperpolarization activated current and resting membrane potential during development (Linear fit (red) slope: -2.84 mV/nS, intercept: -57.67 mV, R^2^=0.71, n=20). **f.** Two- photon microscopy image depicting a proximal and a distal extracellular stimulating location positioned closely to an Alexa-594 filled L2/3 PC dendrite (left), and the resulting EPSPs (proximal – black, distal – green). **g.** EPSP halfwidth decreases with dendritic distance (Linear fit (red) slope: -0.42 ms/µm, intercept: 161.01 ms, R^2^=0.4, n=35). **h.** EPSPs were significantly slower in proximal dendritic locations (147.07 ± 12.87 ms and 85.57 ± 6.83 ms for proximal and distal events, respectively, p=5.31*10^-5^, t(33)=4.64, Student’s two-sample t-test, n=13 and 22, respectively).

Next, we mapped the distance-dependence dendritic EPSP kinetics in 1 week old animals (Fig. 6f). We found a surprising ‘non-Rallian’ negative correlation between EPSP halfwidth and stimulation distance from the soma (Fig. 6g, h), similar to our previous results from mature neurons with HCN channels blocked (Fig. 3a,b). This suggests the presence of nonlinear boosting mechanisms (e.g., NMDA) in proximal dendrites of developing L2/3 PCs, similar to mature cells, but which are unabated due to the lack of HCN at this early developmental stage.

### I_h_ dependent neuromodulation of L2/3 PC activity by serotonergic inputs

Whether the mature brain possesses mechanisms to modulate HCN in L2/3 PCs is unknown. Nonetheless, previous work in cortical L5 PT-type PCs shows that I_h_ currents can be modulated following 5-HT_7_ serotonergic receptor activation(53). L2/3 PCs are extensively innervated by serotonergic axons from the dorsal raphe (54), providing a potential framework to modify HCN availability. Single-cell RNA-seq (29) showed that L2/3 PCs do express 5-HT_7_ receptors, interestingly, at higher levels than all other cortical PCs (Fig. 7a). We pharmacologically evaluated a potential role of serotonergic HCN regulation by bath application of 5-carboxamidotryptamine (5-CT; 30 nM), a specific 5-HT_7_ agonist. 5-CT application significantly reduced evoked firing of L2/3 PCs during extracellular stimulation in L4 (Fig. 7b). Current clamp experiments revealed that the rectification of steady-state voltage responses to hyperpolarizing current injection was amplified with 5-CT (Fig. 7c), likely due to the increased availability of HCN channels due to a rightward shift in their activation voltage-dependence (Fig. 7d). 5-CT also reduced L2/3 input resistance potentially via HCN regulation, however, did not yield a depolarizing effect on the resting membrane potential nor affect AP threshold (Fig. 7e). Importantly, 5-CT had no impact on the firing responses of L2/3 PCs when I_h_ currents were blocked with background Cs^+^ throughout the experiment (Fig. 7f). Furthermore, the dampening of neuronal output was not related to an downregulating effect of 5-HT7 on NMDA receptors (Fig. 7g, (55)). Together these results indicate that neuromodulation represents an endogenous mechanism by which HCN availability can be altered in L2/3 PCs. Further studies on the input-specific effects of neuromodulation on dendritic nonlinearities in L2/3 PCs should thus be considered.

**Fig 7.**
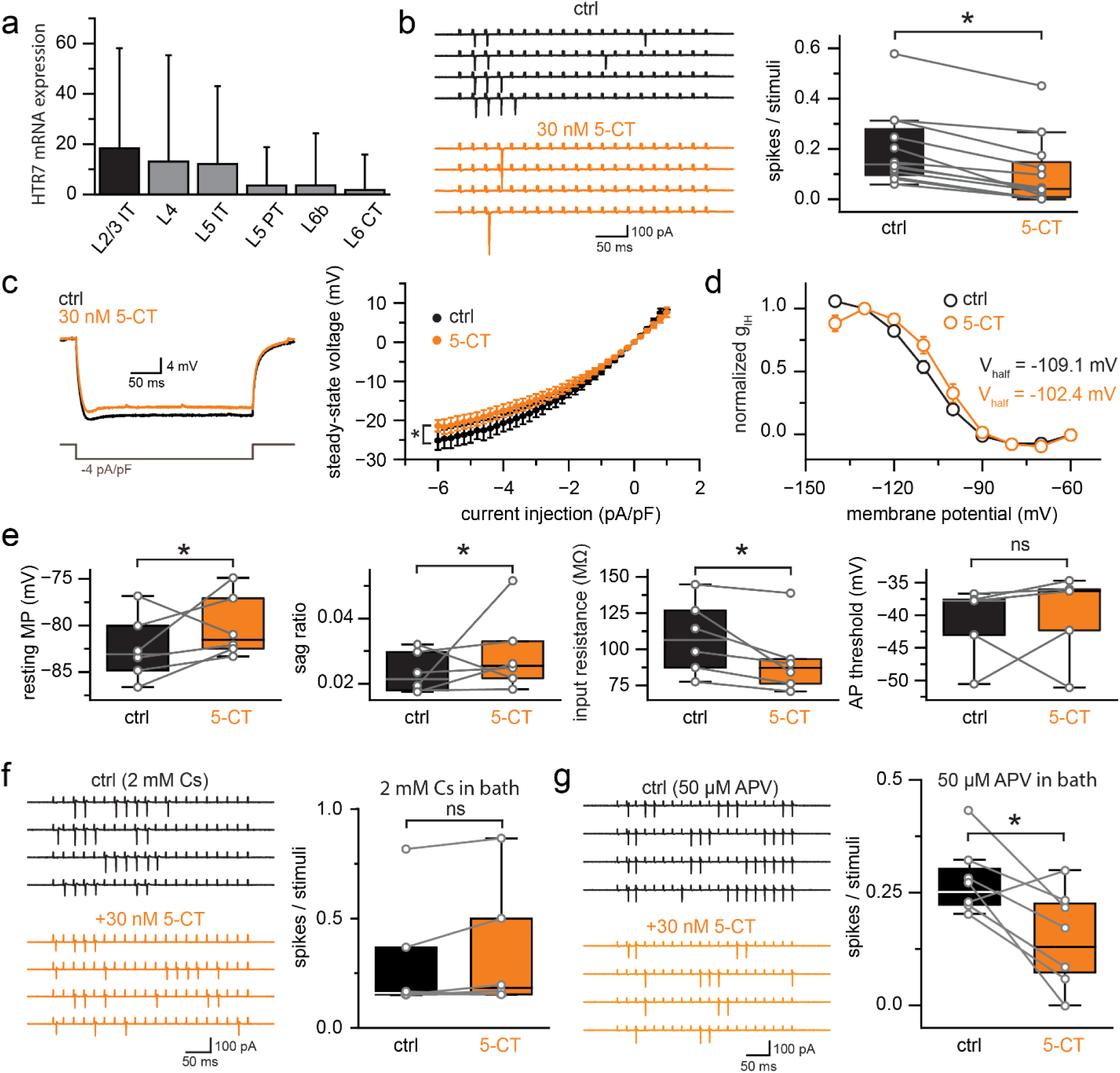
Neuromodulation shifts L2/3 PC excitability via HCN channels. **a.** 5HT_7_ receptor mRNA is abundantly expressed in L2/3 PCs. Data is collected from the online available dataset(29). **b.** Example cell attached recording of a L2/3 PC (left) upon 50 Hz extracellular stimulation in L4 in control conditions (black) and 5-CT bath application (orange). 5-CT significantly reduced L2/3 PC spiking (0.2 ± 0.04 spike/stimuli vs. 0.1 ± 0.04 spike/stimuli for control vs 5-CT application, p=3.57*10^-6^, t(11)=7.92, Student’s paired-sample t-test, n=12 each) **c.** Example current clamp recording of voltage responses to hyperpolarizing current injections (left; control – black, 5-CT – orange). Steady-state voltage response rectification is amplified by 5HT_7_R blockade (right). **d.** 5-CT shifts the voltage dependence of hyperpolarization activated currents (-109.06 ± 0.47 mV vs -102.35 ± 1.09 mV half activation for control vs 5-CT bath application, R^2^ = 0.99 for both). **e.** 5-CT did not alter resting membrane potential (-82.41 ± 1.43 mV vs -80.14 ± 1.38 mV in control vs 5-CT bath application, respectively, p=0.89, t(5)=-1.41, Student’s paired-sample t-test, n=6 each), sag ratio at -2 pA/pF current injection (0.02 ± 2.55*10^-3^ vs 0.03 ± 0.01 in control vs 5-CT bath application, respectively, p=0.82, t(5)=-1.01, Student’s paired-sample t-test, n=6 each) and action potential threshold (-41.13 ± 2.6 mV, -40.08 ± 3.05 mV in control vs 5-CT bath application, respectively, p=0.64, t(4)=-0.4, Student’s paired-sample t-test, n=5 each), but significantly reduced input resistance (108.21± 10.3 mΩ, 92.29 ± 9.9 MΩ in control vs 5-CT bath application, respectively, p=0.02, t(5)=2.75, Student’s paired-sample t-test, n=6 each). **f.** Example recording showing no effect of 5-CT bath application when HCN channels were blocked (left: example recordings, right: 0.3 ± 0.11 spike/stimuli vs 0.34 ± 0.12 spike/stimuli in control vs 5-CT bath application, respectively, p=0.13, t(5)=-1.81, Student’s paired-sample t-test, n=6 each). **g.** NMDA independent 5-CT effect on L2/3 PC excitability (0.27 ± 0.03 spike/stimuli vs 0.14 ± 0.04 spike/stimuli in control vs 5-CT bath application, respectively, p=0.01, t(7)=3.66, Student’s paired-sample t-test, n=7 each).

## DISCUSSION

Precise communication between cortical neurons relies on discrete pre- and postsynaptic forms of information transfer at synapses. The majority of cortical synapses terminate along the dendritic tree (56) where, based on local active and passive electrical properties, various types of nonlinear transformations occur. The spatially and electrically disjointed nature of dendrites (57) from the rest of the cell allows for local, soma-independent computation. For example, perturbing active dendritic elements (i.e., local conductances) can impair several fundamental cortical circuit tasks (36) such as perception (14), underlining the necessity for compartmentalized information processing. Further, single-subunit model simulations are unable to reproduce *in vivo* observed activity patterns of L2/3 PCs(9). Here, by combining *ex vivo* current clamp, synaptic stimulation, and ion channel recordings with empirically informed, biophysically realistic simulations, we demonstrate that L2/3 PCs in sensory cortex express a previously unappreciated, proximally-biased HCN channel distribution. Due to an apparent universal HCN expression in L2/3 cells from each cortical region, this arrangement should allow for modification of synaptic gain across all primary sensory areas of cortex. One of the basic functions of L2/3 PCs in sensory cortex is to encode first-order information, such as the orientation of visual input(2, 58, 59). Several investigations propose that arithmetic summation of a number of similarly tuned synapses will dictate whether these neurons will fire in response to sensory input (60). However, numerous dendritic nonlinearities have been established in L2/3 PCs, such as NMDA receptors (8), sodium channels (11), and calcium channels (10, 61), which also contribute to this process. Here we demonstrate that L2/3 PCs express a functionally relevant population of HCN channels well situated to modify the gain of cortical sensory and contextual information processing.

We report that HCN channels in L2/3 constrain fundamental aspects of cellular excitability, such as input resistance and resting membrane potential. This is likely due to a strong proximal somato-dendritic distribution of HCN channels, as observed in immunohistochemical experiments here. Likely due to their influence on input resistance, HCN blockade resulted in greatly increased AP firing during extracellular stimulation. L2/3 HCN channels strongly dampened excitatory synaptic inputs, however, only to those arriving at proximal dendritic locations. In line with this result, HCN channels reduced the timecourse of synaptic inputs following local stimulation in L4, but left those following L1 stimulation unaffected. Together with additional input-specific pharmacological experiments, NEURON simulations demonstrated that this input-specific synaptic phenomenon could indeed be explained by a proximal HCN channel distribution in dendrites putatively situated in L4 and L2/3. This was apparently only possible with a concurrent proximal NMDA recruitment bias. This predicted global organization should not be confused with observed differences in NMDA recruitment along the axis of individual L2/3 basal dendrites (47). Our predictions of NMDA distributions require further evaluation at the spine-synapse level. Although a proximal-basal dendrite NMDA receptor distribution is in line with findings from L5 PT-type PCs (62), HCN channels are instead enriched distal dendritic compartments in these PC types (19, 31). Thus the unique proximal HCN recruitment in L2/3 PCs permits pathway-specific control of synaptic amplification, as bottom-up (i.e., L4 originating) inputs target proximal dendritic compartments homogeneously (36). This was in opposition to top-down inputs, whose axons largely target distal dendrites in L2/3 PCs (63), as these synapses were unaffected by HCN channel blockade. To fully understand how compartment-specific pathways are functionally regulated, further anatomical and *in vivo* imaging experiments are needed to establish the distribution and synaptic dynamics of L4 and L1 inputs along the L2/3 PC dendritic tree.

HCN channels appear well suited to regulate the synaptic and intrinsic properties of L2/3 PCs, for example, if their availability is plastic or developmentally regulated. Previously, we found that HCN channel availability was upregulated in GABAergic interneurons during development, resulting in reduced propagation of subthreshold signals in mature axons (51). Interestingly, we found that L2/3 PCs in young animals (1 week old) lacked HCN currents. Together, these results suggest a universal developmental HCN upregulation across neuron types. In L2/3 PCs, the HCN channel maturation window was rapid and almost complete by the second postnatal week. Therefore critical period plasticity (64) and developmental I_h_ upregulation occur during a similar developmental window, suggesting a causal relationship. In addition to the robust developmental regulation of HCN channels, we also found that modification of HCN channel availability was also possible in the adult cortex, following serotonergic (5-HT_7_) receptor-mediated biophysical regulation. This was likely due to a metabotropic process that increases cytosolic cAMP levels by stimulating adenylate cyclase(65) which in turn modulates HCN channel properties. This raises the intriguing possibility that neuromodulation could induce nonstationary conditions in mature L2/3 circuits, for example, in re-biasing of bottom-up vs. top-down information, experience-dependent plasticity (66), or even for reopening of critical window-like states (67). A recent finding demonstrated increased spine density and stability during early development due to enhanced 5-HT release in L2/3 PCs of the prefrontal cortex(68), which suggests that serotonin is responsible for not only the morphological, but the physiological regulation of L2/3 PC spines as well.

As L2/3 PCs are one of the most abundant cell populations of the neocortex, HCN channels in these cells may exert overwhelming control over cortical excitability and thus cognition and memory. Cell non-autonomous HCN channel dysregulation manifests in clear behavioral changes, such as altered spatial learning (69) or impaired working memory (70). Curiously, higher HCN1 protein levels in the postmortem brain also correlate with cognitive resilience in aging humans (71). Therapeutic HCN channel modulation aimed at restoring healthy brain function in distinct pathophysiological states has consequently garnered significant interest (72–74), as both HCN1 and HCN2 are linked to neuropathic pain (75) and for treating major depressive disorder (73, 74, 76). Here we demonstrate that activation of the serotonergic receptor 5-HT_7_ alters HCN channel function in L2/3 PCs, resulting in reduced PC excitability, similar to L5 PT-type PCs (53). 5-HT_7_ receptor targeting agents are highly effective psychopharmacological compounds. Agents such as the antidepressant *vortioxetine* has a relevant therapeutic concentration that functionally suppresses 5-HT_7_ (77), and the mood-stabilizing antipsychotic agent *lurasidone* possesses an even higher binding affinity to 5-HT_7_ (78). Commonly used anesthetic drugs also modulate HCN channel action (79, 80), implying the need to investigate anesthesia-related mechanisms and confounds in L2/3 in both therapeutic and experimental settings. Together, we argue that HCN channels are present and functionally relevant in L2/3 PCs. As L2/3 PCs are one of the largest cell populations in the brain, and as HCN channels are the target of several current therapeutics, future research should consider their role in these cells in basic and clinical studies.

## METHODS

### Acute slice preparation

All animal procedures were approved by the Emory University IACUC. Acute slices from cortex were prepared from mature C57Bl/6J mice (6–10 weeks old). Male and female mice were used for all experiments with data collected from ≥3 mice per experimental condition. Mice were first anesthetized with isoflurane and perfused with ice-cold cutting solution (in mM) 87 NaCl, 25 NaHO_3_, 2.5 KCl, 1.25 NaH_2_PO_4_, 2 MgCl_2_, 1 CaCl_2_, 10 glucose, and 75 sucrose. Thereafter, mice were terminated by decapitation and the brain immediately removed by dissection. Brain slices (250 μm) were sectioned in the coronal plane using a vibrating blade microtome (VT1200S, Leica Biosystems) in the same solution. Slices were collected from V1 and from S1, M1 and A1 for a subset of experiments. Slices were transferred to an incubation chamber and maintained at 34°C for ∼30 min and then at 23–24°C thereafter. During whole-cell recordings, slices were continuously perfused with (in mM) 128 NaCl, 26.2 NaHCO_3_, 2.5 KCl, 1 NaH_2_PO_4_, 1.5 CaCl_2_, 1.5MgCl_2,_ and 11 glucose, maintained at 30.0°C ± 0.5°C. All solutions were equilibrated and maintained with carbogen gas (95% O_2_/5% CO_2_) throughout.

### Intracranial viral injections

5-8 week old mice were injected with AAV.CAMKII.C1V1/eYFP (0.2 µl total) in the lateromedial visual area (LM) or the lateral geniculate nucleus (LGN). During intracranial injections mice were head-fixed in a stereotactic platform (David Kopf Instruments) using ear bars, while under isoflurane anesthesia (1.5-2.0%). Thermoregulation was provided by a heating plate using a rectal thermocouple for biofeedback, thus maintaining core body temperature near 37°C. Bupivacaine was subcutaneously injected into the scalp to induce local anesthesia. A small incision was opened 5-10 minutes thereafter and a craniotomy was cut in the skull (< 0.5 μm in diameter) to allow access for the glass microinjection pipette. Coordinates (in mm from Bregma) for microinjection in the LM were X = ± 3.5; Y = -3.87; α = 45°; Z = 0.7, coordinates for the LGN were X = ± 2.0, Y = -2.0; α = 0°; Z = -2.45. Viral solution (titer 1 x 10^13^ vg/mL) was injected slowly (∼0.02 μL min-1) by using a Picospritzer (0.2 μL total). After ejection of virus, the micropipette was held in place (5 min) before withdrawal. The scalp was closed with surgical sutures and Vetbond (3M) tissue adhesive and the animal was allowed to recover under analgesia provided by injection of carprofen and buprenorphine SR. After allowing for onset of expression, animals were sacrificed acute slices were harvested.

### Electrophysiology

L2/3 PCs were targeted for somatic whole-cell recording in the layer 2/3 region of primary visual cortex by gradient-contrast video microscopy. Electrophysiological recordings were obtained using Multiclamp 700B amplifiers (Molecular Devices). Signals were filtered at 6–10 kHz and sampled at 50 kHz with the Digidata 1440B digitizer (Molecular Devices). For whole-cell recordings, borosilicate patch pipettes (2.5-3.5 MΩ tip resistance) were filled with an intracellular solution containing (in mM) 124 potassium gluconate, 2 KCl, 9 HEPES, 4 MgCl_2_, 4 NaATP, 3 l-ascorbic acid, and 0.5 NaGTP. Pipette capacitance was neutralized in all recordings and electrode series resistance was compensated using bridge balance in current clamp. Liquid junction potentials were uncorrected.

For cell attached recordings seal resistance of >1GΩ was established, for accurate measurement of action potentials. Cells were held at -80 mV. A stimulating electrode (2.5-3.5 MΩ tip resistance, filled with extracellular solution) was placed in either layer 4 or layer 1 region of the visual cortex, at least 200 µm away from the recorded soma. Stimulation intensities were calibrated to reliably elicit at least 3, but no more than 10 action potentials during the 20 consequent stimuli (50 Hz, 30 s inter-trace intervals, 10-50 µs stimulus length). Stability of the elicited responses was confirmed by an initial 15 repeated measurements. Recordings were post hoc filtered at 2 kHz to remove stimulation artifacts.

Recordings had a series resistance <20 MΩ. Membrane potentials were maintained near –80 mV during current-clamp recordings using constant current bias. AP trains were initiated by somatic current injection (300 ms) normalized to the cellular capacitance in each recording measured immediately in voltage clamp after breakthrough (81). Cellular electrical and firing parameters were established by recording voltage responses to current steps between -6 pA/pF and 20 pA/pF. Sag potential was measured at the -6 pA/pF step, by comparing the negative peak within the first 100 ms of the hyperpolarized voltage response and the mean voltage during the last 50 ms of the stimuli. Input resistance was measured by averaging voltage responses to a 20 pA prepulse current step. Sag ratio was determined as the ratio between the sag amplitude and the steady-state response at the -6 pA/pF current step, except when cells were recorded for measuring neuromodulation (Fig 7.), as those experiments were significantly more time-consuming. In these cases, -2 pA/pF current injection was provided.

To establish distance dependent EPSP kinetics, cells were filled with 10 µM Alexa-594. Dendrites were visualized after cell fill with Alexa 594 (10μM) with two-photon laser scanning microscopy using a commercial scan head (Ultima; Bruker Corp) fitted with galvanometer mirrors using a 60x, 1.0 NA objective. The fluorophore was allowed to equilibrate for at least 10 minutes before imaging. The stimulating electrode, filled with extracellular recording solution, was placed in close proximity (<5 µm) of the two-photon visualized dendrite. Stimulation intensity was calibrated to elicit an EPSP with <1 ms delay and approximately 50% failure rate (54.21 ± 8.72%, n= 41). The 50% failure rate was necessary in order to maximize the possibility of stimulating a single input. For L1, L4-Type I, and L4-Type II input stimulations the procedure was the same. Synaptic summation was probed by eliciting 5 consequent responses at 50 Hz. Stimulation intensity was elevated during these recordings to elicit synaptic responses every time for the first stimuli as failures introduced by lower stimulation intensities can yield inconsistent input activation during the 5 stimuli. Inhibitory blocker SR95531 (10 µM) was only included during signal summation (50 Hz stim) experiments, as our intention was to examine single inputs in an intact system. However, repeated stimuli recruited significant inhibitory components as well, therefore the use of inhibitory blockers was warranted for the uncontaminated analyses of our recordings for the 50Hz repetitive stimulation experiments. Z-stacks were created from the stimulated dendritic region and the connected soma as well for 3D reconstruction in order to establish exact dendritic distances. The group of distal dendritic branches only included a fraction of apical dendrites, however proximal dendrites included both a fraction of apical and basal dendrites, for two reasons; first, basal dendrites are significantly shorter than apical dendrites and our recordings did not include any basal dendrites extending beyond 200 µm, and second, in superficial L2 pyramidal cells classifying dendrites that extend radially into apical or basal classes in ambiguous.

For voltage-clamp recordings, cells were patched in normal extracellular solution to first record their firing patterns. Voltage protocols for I_h_ current measurements consisted of 300 or 500 ms long voltage steps between −150 and −60 mV. Activation time constant was measured by fitting the initial 100 ms of the current response with a single exponential function. In a subset of recordings CsMeSO4 (2 mM) was bath applied after recording control current responses, to establish cesium-sensitive I_h_ current kinetics and voltage dependence by subtraction. Series resistances were between 6–20 MΩ (75–80% compensated with 53 µs lag) and were constantly monitored. Data were discarded if the series resistance changed more than 25%.

The dynamic clamp system was built in-house based on a previous publication(82), related online available materials available here: (http://www.dynamicclamp.com/). The equations governing the implemented gHCN were identical to those used in the NEURON model construction. Synaptic conductances were built-in predefined conductances available from http://www.dynamicclamp.com/.

### Intracellular solution perfusion

Intracellular solution was perfused by a custom-built tubing and relay system attached to the pipette holder. Briefly, the pipette was filled with normal intracellular solutions to measure firing patterns and subsequent control measurements of spontaneous and elicited EPSPs. Next, a second intracellular solution was introduced to the pipette through a small tube drilled through the pipette holder, by applying miniscule positive pressure. The intracellular fluid levels were marked before the recording and through a second tube the intracellular solution was siphoned out, to not alter established pipette capacitance. During initial trial experiments using fluorophores in the second intracellular solution it was determined that two minutes is enough for complete exchange of the intracellular solution and intracellular equilibration as well. Data were discarded if the series resistance or pipette capacitance changed more than 20%.

### Immunohistochemistry

Two-month-old mice were anesthetized with Nembutal (50mg/mL, i.p.) and perfused transcardially with 4% paraformaldehyde (PFA), 0.05% glutaraldehyde in phosphate buffered saline (PBS). After perfusion, brains were removed from the skull and immersed in 4% PFA overnight at 4°C. Coronal 100 μm-thick sections were then cut on a Vibratome (Leica V1000). Brain slices were permeabilized and incubated in blocking buffer for one hour at room temperature. After blocking, sections were overlaid with primary antibody to HCN1 (1:200, Thermo Fisher Scientific) overnight at 4 °C, then subjected to a 1 h incubation with secondary antibody (Goat anti-Rabbit IgG, 1:200, Thermo Fisher Scientific) at room temperature. Stained sections were mounted with DAPI-containing mounting solution and sealed with glass coverslips. Confocal imaging was carried out on a Leica SP5 MP.

### NEURON modeling

Computer simulations were performed using the NEURON simulation environment (versions 7.5 and 7.6, downloaded from http://neuron.yale.edu). A constrained NEURON model with an active somatic compartment was downloaded from the publicly available Cell Type Database of the Allen Institute (http://celltypes.brain-map.org/experiment/electrophysiology/487661754). Passive properties (axial resistance, specific membrane capacitance and leak conductance density) were altered manually to recreate passive membrane potential responses of whole-cell recorded L2/3 PCs for given stimulus intensities. Both transient and persistent potassium channels were taken out of the model to gain simulation speed as the aim of the simulations was to reproduce subthreshold EPSP dynamics. Final simulations were rerun with intact sodium currents as well to confirm the lack of alterations introduced by these channels. The I_h_ conductance was based on the built-in Hodgkin–Huxley model of NEURON with freely adjustable sets of parameters(83). I_h_ model parameters were constrained by our whole-cell voltage clamp recordings. The steady-state activation was governed by the following equation:

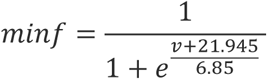

where v is the local membrane potential. Due to the lack of apparent inactivation, the model operated with a single gate (activation).

The activation and deactivation time constant was defined as

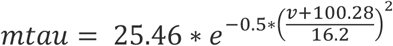

To simulate EPSP kinetics, a movable spine (length: 0.6 µm, diameter: 0.6 µm, axial resistance: 150 Ωcm), with spine neck (length: 2 µm, diameter: 0.1 µm, axial resistance: 100 Ωcm) was created. Leak conductance was inserted into both compartments, along with the fitted conductance model. Synaptic inputs were supplemented by using NEURON’s built-in AlphaSynapse class. To probe the contribution of specific dendritic conductances to shaping the distance-EPSP halfwidth relationship, selected conductance models were inserted into both apical and basal dendrites with a Gaussian distribution. This curve was selected as modulation of the Gaussian curve’s width and y-offset can reproduce a wide variety of feasible distributions such as linear, exponential, or hot-zone-like expression. To model the contribution of dendritic sodium, potassium, and calcium channels to input modulation, NaTs, Kv3_1 and Ca_LVA models were selected. Distribution approximation was done by grid search to avoid local fitting minimums. NMDA receptors were implemented based on a previous publication (84). During the fitting procedure, NMDA activation voltage and AMPA:NMDA ratios were adjusted by a grid search algorithm.

### Statistics

Averages of multiple measurements are presented as mean ± SD. Data were statistically analyzed by ANOVA test using Origin software and custom-written Python scripts. Normality of the data was analyzed with Shapira-Wilks test.

## Data and software availability

All code used for simulating single and multicompartmental NEURON models will be made available on GitHub and public repositories upon VOR.

---

**Supplementary Figure 1.**
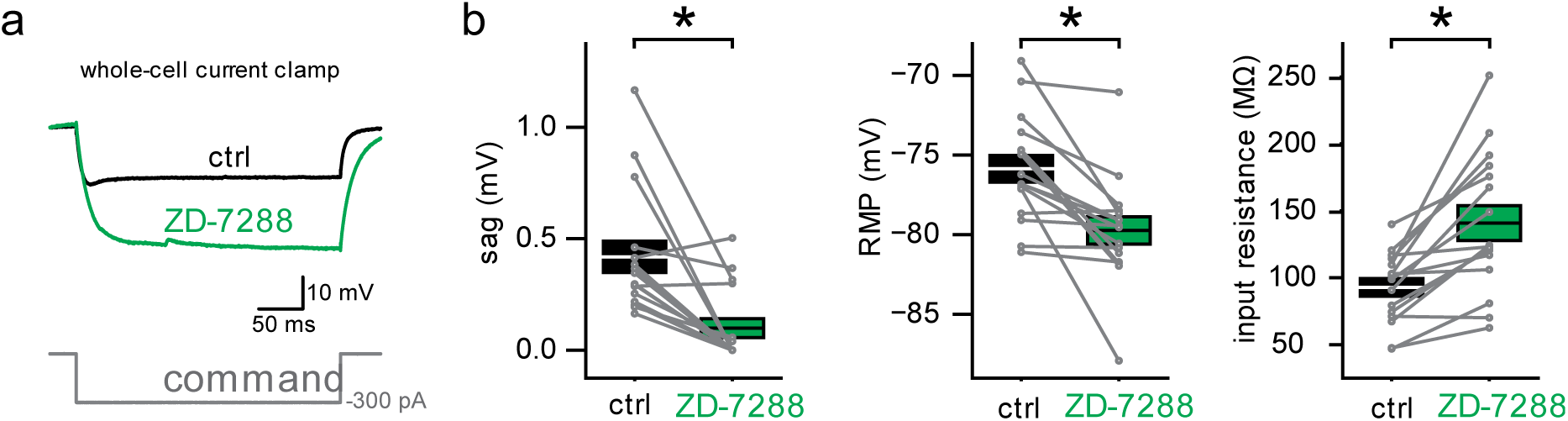
ZD-7288 application results in similar active and passive electrical alterations. **a.** Representative effect of ZD-7288 on voltage responses to hyperpolarizing current injections. **b.** ZD-7288 abolished sag voltage (0.42 ± 0.07 mV vs 0.1 ± 0.04 mV in control vs ZD-7288 application, respectively, p=4.5*10^-4^, t(15)=4.47, Student’s paired-sample t-test, n=16), decreased the resting membrane potential (-75.86 ± 0.84 mV vs -79.73 ± 0.86 mV in control vs ZD-7288 application, respectively, p=4.41*10^-4^, t(15)=4.48, Student’s paired-sample t-test, n=16), and increased the input resistance (92.74 ± 6.59 MΩ vs 140.8 ± 13.07 MΩ in control vs ZD-7288 application, respectively, p=1.79*10^-4^, t(15)=-4.93, Student’s paired-sample t-test, n=16).

**Supplementary Figure 2.**
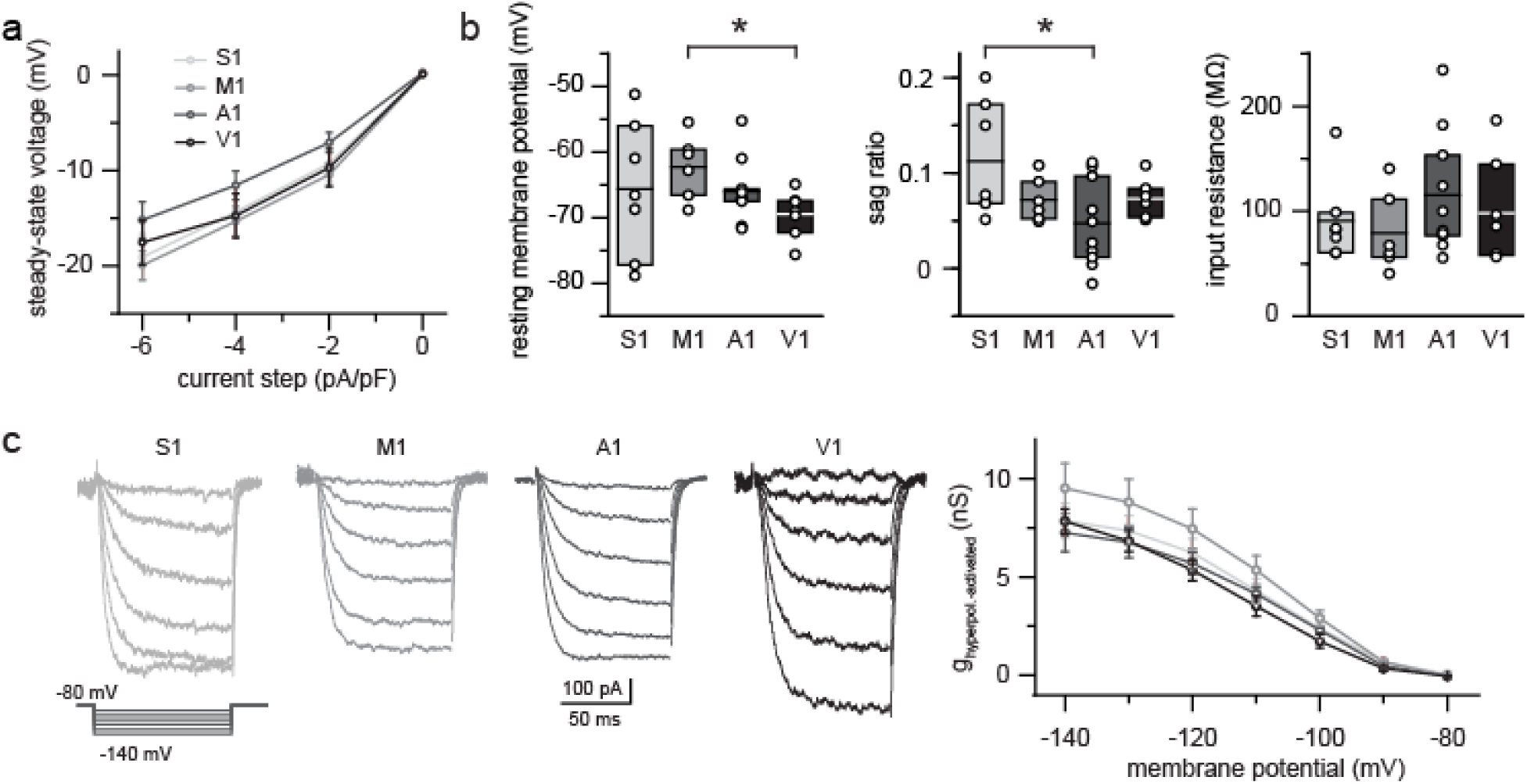
Similar Ih current and effect in different cortical areas. **a.** Steady-state rectification of L2/3 PCs in the primary somatosensory cortex (S1), primary motor cortex (M1), primary auditory cortex (A1) and primary visual cortex (V1). **b.** Similar resting membrane potential (left), sag ratio (middle) and input resistance (right) of L2/3 PCs in different cortical areas. **c**. Example hyperpolarization activated current recordings from four cortical areas (left), with similar voltage dependent activation (right).

**Supplementary Figure 3.**
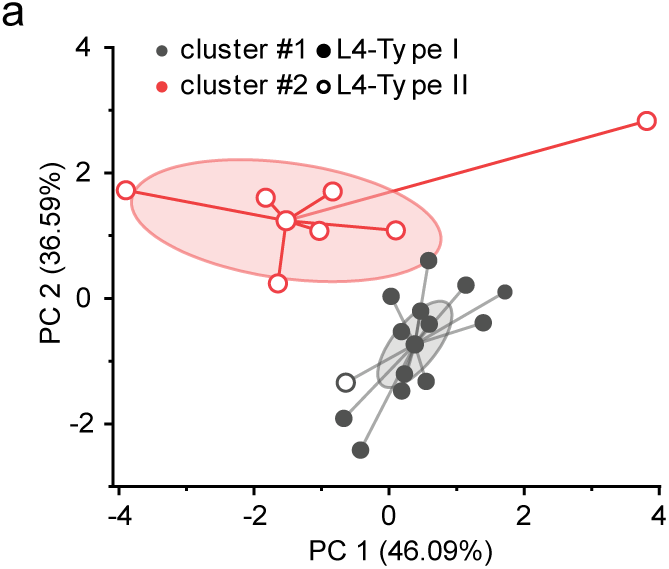
L4-Type I and L4-Type II inputs are well separated by k-means clustering. k-means clustering results plotted along the first and second principal component. Ellipses denote the confidence interval; edges are connected to the cluster center. Hollow circles represent L4 Type II inputs and filled circles represent L4-Type I inputs determined by the experimenter.

**Supplementary Figure 4.**
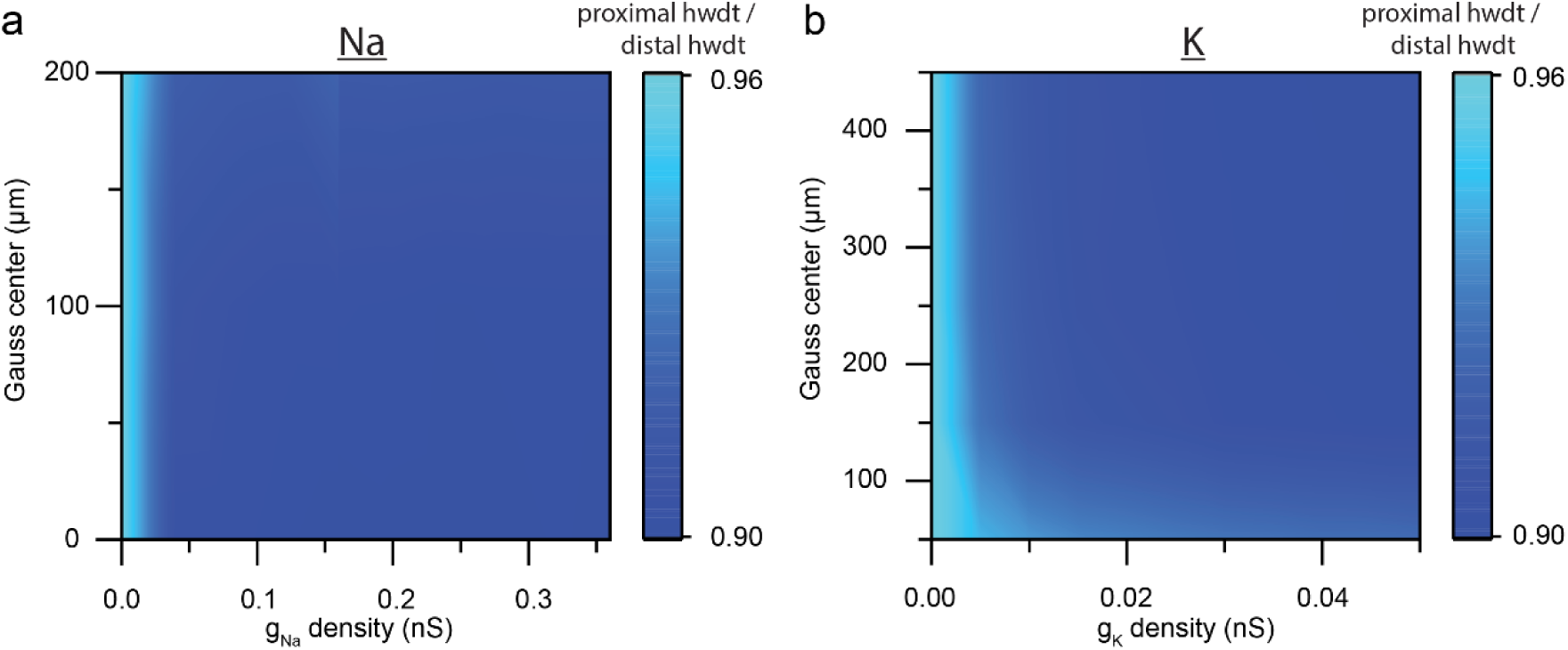
The presence of dendritic sodium and potassium channels cannot explain wider proximal synaptic inputs. **a.** Sodium channels cannot explain the distance-EPSP halfwidth relationship observed when I_h_ was blocked. Color coding denotes the ratio between proximal and distal EPSP halfwidths. Notice the miniscule range of the color coding. **C.** Same as panel **b.** but for voltage dependent potassium channels.

**Supplementary Figure 5.**
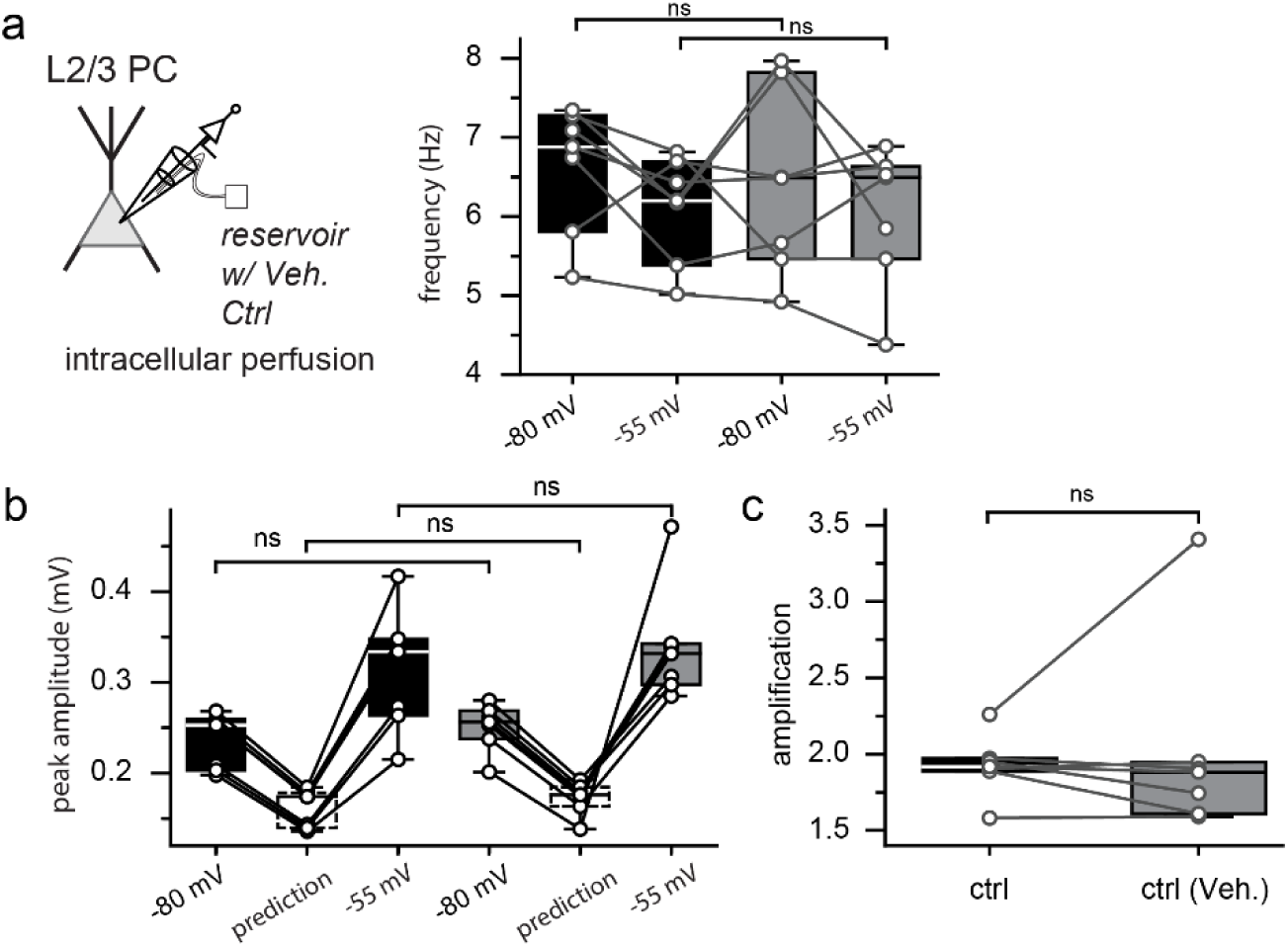
Pipette perfusion does not alter spontaneous circuit activity measurements. **a.** sEPSP frequency is unaltered by pipette perfusion (6.63 ± 0.3 and 6.4 ± 0.44 for ctrl -80 mV and perfused ctrl -80 mV, p=0.57, t(6) = 0.59, 6.1 ± 0.25 and 6.03 ± 0.33 for ctrl -55 mV and perfused ctrl -55 mV, p=0.83, t(6) = 0.23, n=7 for each, Student’s paired-sample t-test). **b.** sEPSP peak amplitudes are stable during pipette perfusion experiments (0.23 ± 0.01 and 0.25 ± 0.01 for ctrl -80 mV and perfused ctrl -80 mV, p=0.43, t(6) = -0.83, 0.16 ± 0.01 and 0.17 ± 0.01 for ctrl prediction and perfused prediction, p=0.43, t(6) =-0.83, 0.31 ± 0.03 and 0.34 ± 0.02 for ctrl -55 mV and perfused ctrl -55 mV, p=0.24, t(6) = -1.31, n=7 each, Student’s paired-sample t-test). **c.** Pipette perfusion does not alter measured synaptic amplification (1.92 ± 0.07 and 2.01 ± 0.24 for ctrl and perfused ctrl, p=0.65, t(6) = -0.48, n=7 each, Student’s paired-sample t-test).

**Supplementary Figure 6.**
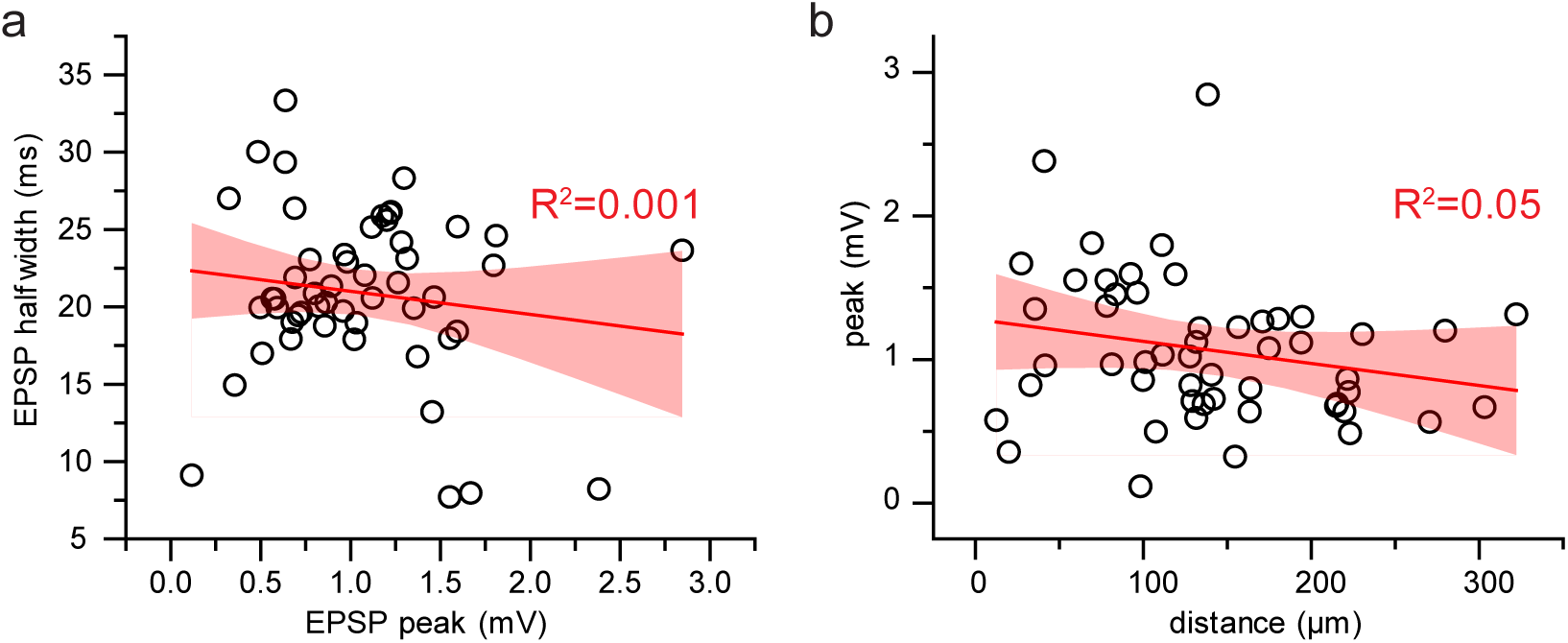
Elicited EPSP halfwidth in not dependent on event amplitude. **a.** EPSPs elicited by electrical stimulation show no correlation between event amplitude and halfwidth (R^2^= 0.021, n=53). **b.** EPSP peak is not dependent on dendritic distance from the soma (R^2^= 0.021, n=53).

